# Characterizing neuroanatomic heterogeneity in people with and without ADHD based on subcortical brain volumes

**DOI:** 10.1101/868414

**Authors:** Ting Li, Daan van Rooij, Nina Roth Mota, Jan K. Buitelaar, the ENIGMA ADHD Working Group, Martine Hoogman, Alejandro Arias Vasquez, Barbara Franke

## Abstract

**Background:** Attention-Deficit/Hyperactivity Disorder (ADHD) is a prevalent neurodevelopmental disorder in children and adults. Neuroanatomic heterogeneity limits our understanding of the etiology of ADHD. This study aimed to parse neuroanatomic heterogeneity of ADHD, and to determine whether subgroups could be discerned in patients based on subcortical volumes.

**Methods:** Using the dataset from the ENIGMA-ADHD Working Group, we applied exploratory factor analysis (EFA) to subcortical volumes of 993 boys with and without ADHD, and to subsamples of 653 adult men, 400 girls, and 447 women. Factor scores derived from the EFA were used to build networks. A community detection (CD) algorithm clustered participants into subgroups based on the networks.

**Results:** Three factors (basal ganglia, limbic system, and thalamus) were found in boys and men with and without ADHD. The factor structures for girls and women differed from those in males. Given sample size considerations, we concentrated subsequent analyses on males. Male participants could be separated into four communities, though Community 3 was absent in healthy men. Significantly case-control differences of subcortical volumes were observed within communities in boys with increased effect sizes, but not in men. While we found no significant differences in ADHD symptom severity between communities in boys or men; affected men in Community 1 and 4 presented comorbidities more frequently than those in other communities.

**Conclusion:** Our results indicate that neuroanatomic heterogeneity in subcortical volumes exists, irrespective of ADHD diagnosis. Effect sizes of case-control differences appear more pronounced at least in some of the subgroups.

## Introduction

Attention-Deficit/Hyperactivity Disorder (ADHD) is a prevalent neurodevelopmental disorder characterized by age-inappropriate inattention (IA) and/or hyperactivity and impulsivity (HI) (1). ADHD frequently persists from childhood into adulthood, with a prevalence of 3.4-5.3% in childhood/adolescence and 2.5% in adulthood (2-4).

ADHD is a heterogeneous disorder on the clinical, cognitive, genetic, and neuroanatomic level. Clinically, there is strong interindividual variation in psychiatric and somatic comorbidities across the lifespan (5). Most individuals with ADHD have deficits in one or more cognitive domains, but there is substantial overlap between ADHD and controls (6-8). The estimated heritability of ADHD is 70-80%; and common genetic variants with small effect size are the major contributors to genetic susceptibility to ADHD (9). Considerable heterogeneity is also present in structural and functional brain architecture. The most consistent findings were observed for structural brain alterations in subcortical regions (10). To overcome the limitations of small sample size studies, the ENIGMA-ADHD Working Group conducted a large mega-analysis (1713 cases and 1529 controls) across the lifespan (11). This analysis confirmed earlier findings of reduced caudate nucleus, putamen, and total intracranial volumes in ADHD, and identified smaller nucleus accumbens and amygdala volumes in individuals with ADHD compared with healthy controls. Volumetric case-control differences were most prominent in childhood. However, the effect sizes were small, possibly reflecting neurobiological heterogeneity of ADHD.

Classification methods have been used to investigate heterogeneity within groups (12). In ADHD research, community detection (CD), a graph-theoretical measure, has been applied to identify clusters of children with different neuropsychological performance profiles across a battery of tasks (13). A similar method was used to identify three subgroups of children with ADHD presented distinct profiles of emotional functioning associated with clinical outcome (14). Taximetrics analysis was applied in a sample of adolescents with ADHD, resulting in three subgroups with different profiles of executive functioning and motor inhibition (15). In combination with other studies on the heterogeneity of functional brain architecture in ADHD (16, 17), the results of these investigations suggested that differences in clinical and neurobiological presentation and course of ADHD may be captured in distinct subpopulations. Moreover, while cases and healthy controls were present in the same subgroups, affected individuals within a subgroup were more impaired (8, 13, 15).

As CD methods have been widely applied to brain networks (18), in the current study, we utilized this approach to parse neuroanatomic heterogeneity in ADHD using the subcortical brain volume data from the ENIGMA-ADHD Working Group (n=2493 in total). Our objectives were 1) to examine whether subgroups of participants could be defined based on subcortical volumes and whether this categorization was related to the clinical presentation of ADHD, and 2) to explore whether the effect size of case-control differences would be increased within a subgroup.

## Method

### Participants and ADHD assessment

For the present study, we used available magnetic resonance imaging (MRI) data from the international ENIGMA-ADHD Working Group (http://enigma.ini.usc.edu/ongoing/enigma-adhd-working-group/). The group shares structural MRI scans from children and adults with ADHD, as well as phenotypic information, including comorbidities, IQ, age, and gender from over 35 cohorts across the world. With a rolling inclusion design, new cohorts can join the group at any time, but data freezes are set for each analysis. Each site verified the diagnosis of ADHD and assessment of comorbidities (Table S1). All participating sites had approval from local ethics committees.

To constrain heterogeneity in the ENIGMA-ADHD dataset, we stratified our sample by age and sex. Our subsamples comprised 993 boys (aged 4-14 years), 400 girls (aged 4-14 years), 653 adult men, and 447 women (aged >22 years) (Table 1). We started by applying EFA and CD to the subsample of boys, which was the largest subsample within the dataset. The same method was subsequently applied to the other three subsamples to investigate whether similar subgroups exist in these subsamples.

**Table 1:** Characteristics of participants.

| **Variables** | **Boys** |  | **Girls** |  | **Men** |  | **Women** |  |
| --- | --- | --- | --- | --- | --- | --- | --- | --- |
|  | **Patients** | **Controls** | **Patients** | **Controls** | **Patients** | **Controls** | **Patients** | **Controls** |
| **N** | 563 | 430 | 135 | 265 | 412 | 241 | 223 | 224 |
| **Mean Age** | 11.0 | 10.8 | 10.3 | 10.4 | 32.1 | 32.2 | 36.1 | 35.7 |
| **(SD)** | (1.9) | (1.9) | (1.9) | (1.8) | (8.9) | (8.7) | (10.1) | (11.0) |
| **Mean IQ** | 103.2 | 110.4 | 103.1 | 112.5 | 108.4 | 113.3 | 108.5 | 110.5 |
| **(SD)** | (15.8) | (14.8) | (14.8) | (13.8) | (14.3) | (14.8) | (15.4) | (15.3) |

*Note:* SD: Standard deviation

### Neuroimaging

Structural T1-weighted brain MRI data were collected at each site. All scans were subsequently analyzed using the standardized ENIGMA protocols based on FreeSurfer version 5.1 or 5.3. For each participant, we computed left and right volumes of the nucleus accumbens, putamen, pallidum, caudate nucleus, thalamus, amygdala, and hippocampus, as well as intracranial volume (ICV). For all analyses, we used the mean of the left and right subcortical volume. Outliers were identified as above or below three times the interquartile range, and participants with missing data were excluded from the analysis.

### Exploratory Factor Analysis (EFA)

Exploratory factor analysis (EFA) was applied to reduce the space of subcortical volume data by modeling latent factors, which in general requires 300 cases per analysis (19). In considering non-linear patterns of subcortical brain volumes across age, each subcortical volume was regressed individually with age, age^2, sex, ICV, and sampling site; this was done for children and adults separately. Residuals were used to construct covariance matrices. Squared multiple correlations were built as prior communality estimates. A maximum likelihood method and oblique rotation were used to extract factors. The number of eigenvectors extracted was based on the scree-plot. A variable was considered to load on one factor if the loading on the factor was 0.40 or more. Model fitness was evaluated based on Tucker-Lewis Index (TLI), Akaike information criterion (AIC), and the root mean square error of approximation (RMSEA). The analyses were performed using the psych package in R programming v3.1.1.

### Community Detection (CD)

We applied CD to identify distinct communities of participants based on factor scores generated by the EFA of subcortical volumes. Applying a modularity algorithm, CD identifies clusters of individuals in a network by requiring strong correlation among them (18). CD was performed in three steps. First, n × n weighted, undirected networks were created by correlating participants with each other on their normalized factor scores to provide distance information between subject pairs. For this, a threshold of r = 0.5 was chosen, where reachability remained equal to 1. Subsequently, a weight-conserving modularity algorithm was applied to identify distinct communities of participants in each network (13, 20). To obtain the most optimal partitioning of the network, this algorithm iteratively sorts nodes (participants in this study) into communities until the modularity (Q) reaches a maximum. Q is the number of edges (correlations between participants) falling within communities minus the expected number in a random network. Q ranges between −1 and 1, with positive values indicating that the strength of edges within communities is larger than expected at random.

To assess robustness of the community structure, we examined variation of information (VOI). Briefly, a proportion of edges of a network was randomly perturbed. VOI was calculated as the variance between the original and perturbed networks over a range of alpha, which ranges between 0 and 1 (21).

All CD analyses were performed in Matlab (Mathworks) and the functions provided by Olaf Sporns, Mikail Rubinov, and collaborators (20).

### Statistical Analyses

Age was compared between patients and controls using an independent-samples t-test; Estimated IQ scores were compared between groups with Analysis of Variance (ANOVA) after regressing the effects of age, IQ assessment instrument, and sampling site. For each community, we compared subcortical factor scores between cases and controls using t-tests. Starting with the subsample of boys, we also investigated whether ADHD symptom severity differed among the communities using t-test or ANOVA; Chi-square tests were used to compare the presence of comorbidities between communities. False discovery rate (FDR) was used to correct for multiple comparisons. All analyses were performed in IBM SPSS Statistics 22.

## Results

### Participant characteristics

Demographics of this sample are described in Table 1. Mean age did not differ between cases and controls for boys (t=-1.7, p=0.09), girls (t=0.14, *p*=0.89), men (t=0.22, *p*=0.83), and women (t=-0.37, *p*=0.71). Differences in IQ scores between cases and controls were significant in boys (F=16.7, df=4, *p*=3.8e-13), girls (F=8.8, df=4, *p*=8.6e-7), men (F=5.1, df=4, *p*=5.0e-4), and women (F=3.8, df=4, *p*=0.005).

### EFA on subcortical volumes

Aiming to limit heterogeneity and maximize power, we started with the largest subsample available, which was for boys, and performed EFA on residualized subcortical brain volumes. From the covariance matrix, we extracted three eigenvectors (Figure 1). Volumes of caudate nucleus, globus pallidus, nucleus accumbens, and putamen loaded on the first factor. We interpreted this first factor as “basal ganglia”. The second factor included hippocampus and amygdala, and was interpreted as “limbic system”. The third factor comprised only the thalamus. The three factors accounted for 25%, 16%, and 12% of the total shared variance, respectively.

**Figure 1:**
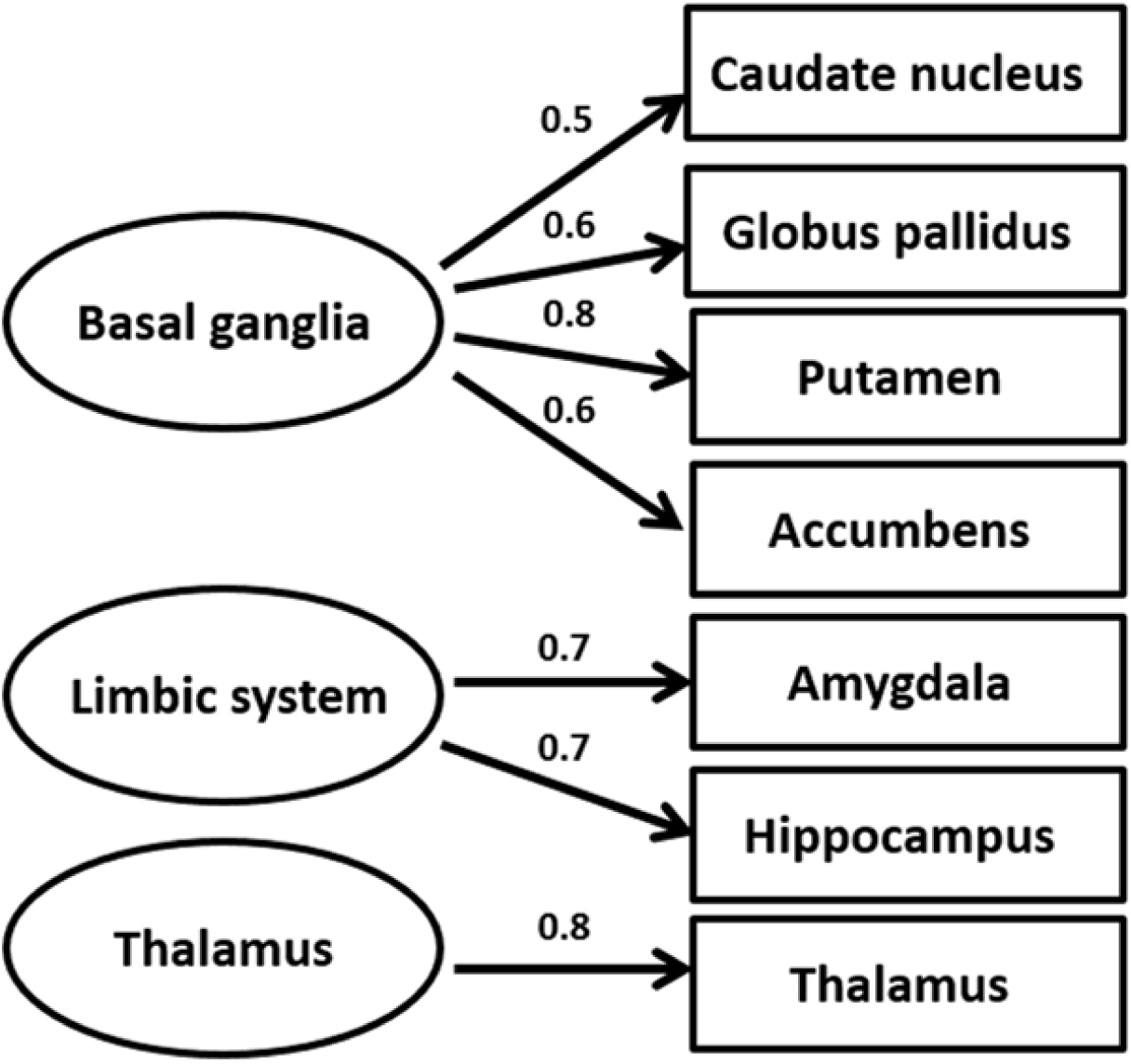
The three-factor model that was generated by EFA in the boys with estimated factor loadings of the latent factors. *Note*: *Similar factor models were generated in boys with and without ADHD separately*.

We next performed EFA in girls (Figure S1). Volumes of caudate nucleus, nucleus accumbens, and putamen loaded on the first factor; the second factor only included the globus pallidus; the third factor comprised by the hippocampus, amygdala, and thalamus volume. The three factors accounted for 16%, 18%, and 20% of the total shared variance, respectively. The comparison of model fitness indicated that this factor structure (TLI=0.77, AIC=39053, RMSEA=0.12) was superior to the one generated in boys (TLI=0.69, AIC=39080, RMSEA=0.14; chi square difference=26.4, *p*=2.2e-16).

EFA was also run for adult men and women, separately. In men with and without ADHD, the same three eigenvectors as in boys were extracted, which accounted for 23%, 17%, and 17%, respectively, of the total shared variance. In women, three eigenvectors were also found, but the factor structure differed from the others (Figure S2). Volumes of nucleus accumbens and putamen loaded on the first factor. The second factor included caudate nucleus, globus pallidus, and thalamus. The third factor comprised hippocampus and amygdala volume. The three factors accounted for 17%, 18%, and 23% of the total shared variance, respectively. Additional model comparison indicated that this factor structure was superior in the subsample of women (TLI=0.06, AIC=44454, RMSEA=0.29) to the factor structure we usually got in subsamples of males (TLI=-0.01, AIC=44518, RMSEA=0.30; chi square difference=66.6, *p*=3.4e-16).

### CD on factor scores of subcortical volumes in boys and men

Given that factor structures differed between males and females, subsequent CD results would have been incomparable between them. For the subsequent CD analyses, we therefore focused exclusively on boys and adult men, where sample sizes were most appropriate for CD-type analyses.

In all boys (with and without ADHD), we observed four distinct communities, each comprising 20-30% of the sample (Figure 2; Table 2). Community 1 was characterized by increased volume in basal ganglia, normal volume in limbic system, and smaller volume of thalamus compared to the average volume of the whole sample. Community 2 showed opposite characteristics for basal ganglia and thalamus to Community 1. Community 3 had smaller basal ganglia and thalamus and larger volume in the limbic system, whereas Community 4 showed the reverse pattern compared to Community 3. Repeating the analysis in boys with and without ADHD separately resulted in largely similar findings (Figure 2).

**Table 2:** The distribution of participants in subsamples in communities.

| **Subsamples** | **Total** | **Patients** | **Controls** |
| --- | --- | --- | --- |
| **Boys (N)** | **992** | **563** | **430** |
| **Community 1** | 220 (22.2%) | 119 (21.1%) | 130 (30.3%) |
| **Community 2** | 270 (27.2%) | 167 (29.7%) | 95 (22.1%) |
| **Community 3** | 234 (23.6%) | 122 (21.8%) | 103 (24.0%) |
| **Community 4** | 268 (27.0%) | 154 (27.4%) | 101(23.5%) |
| **Q values** | 0.45 | 0.45 | 0.46 |
| **Men (N)** | **653** | **412** | **241** |
| **Community 1** | 201 (30.8%) | 127 (30.8%) | 102 (42.3%) |
| **Community 2** | 166 (25.4%) | 90 (21.8%) | 70 (29.0%) |
| **Community 3** | 97 (14.9%) | 79 (19.2%) | 0 |
| **Community 4** | 189 (28.9%) | 116 (28.2%) | 69 (28.6%) |
| **Q values** | 0.47 | 0.46 | 0.49 |

**Figure 2:**
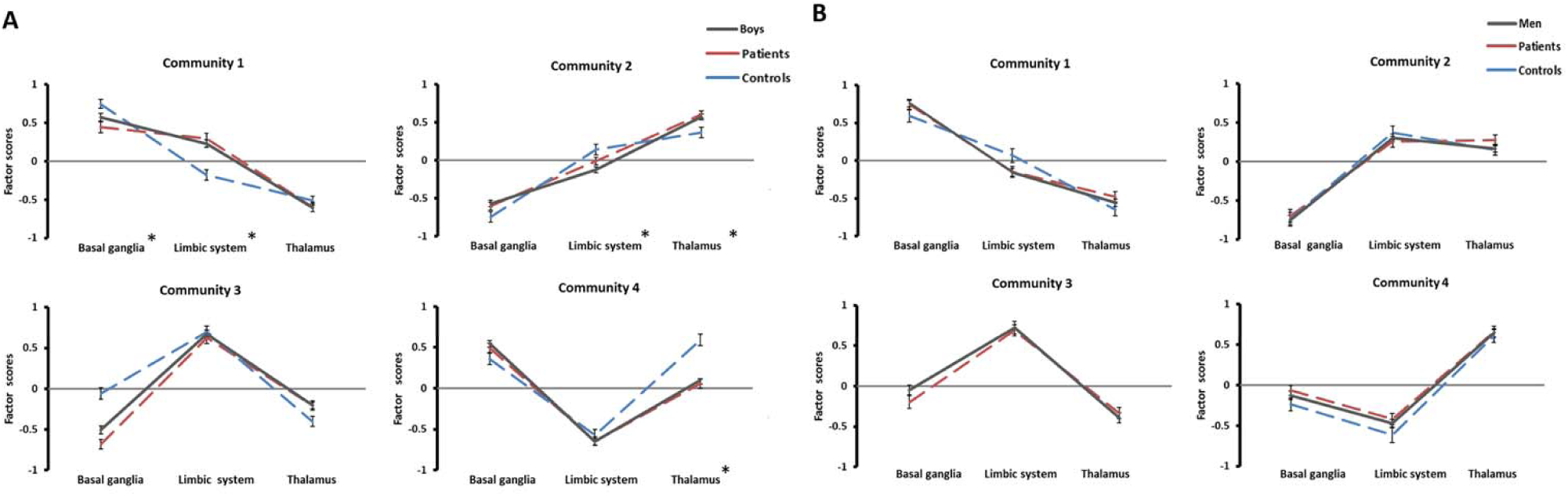
Communities generated by CD. A: Communities in boys; B: Communities in men. *Note: Lines represent participants in each community from CD. Y-axis indicates the mean factor scores for each factor. Error bars: standard error of the mean. * indicates the difference of factor scores between patients and controls are significant*.

Quality control measures, i.e. the quality index (Table 2) and VOI (Figure S3), showed that these communities were significantly different from subgroups generated from random networks, and the networks were robust against chance variation. Furthermore, although the distribution of cases and controls across communities differed among cohorts (Table S2), leave-one-out analyses of the five largest cohorts showed no evidence for specific cohorts driving the community structure, and the same four communities were found in each analysis.

CD in adult men (with and without ADHD) resulted in four communities similar to those observed in boys, each accounting for 15-31% of the sample (Figure 2; Table 2). Cases were distributed across all four communities, but the controls were only present in three communities, with no healthy men in Community 3. The distribution of cases and controls over communities is shown in Table S3 for each cohort.

### Comparison of subcortical factor scores between patients and controls in each community

Within each of the four unique communities observed in boys and men, we investigated whether subjects with and without ADHD showed different volumes in structures contributing to the subcortical factors (Table 3, Figure 2). Boys with ADHD in Community 1 and Community 3 had reduced subcortical volumes in basal ganglia compared to controls; boys with ADHD in Community 1 also had larger volumes in the limbic system than controls. Those with ADHD in Community 2 had smaller volumes in this system than controls. Boys with ADHD in Community 2 and Community 3 also showed larger volumes for thalamus, and those in Community 4 had smaller thalamus volume. Effect sizes for boys ranged from *d*= −0.90 (95% CIs [−1.17, −0.62]) to *d*= 0.65 (95% CIs [0.39, 0.90]) (Table 3). In Supplementary Table S4, we present case-control comparisons for each individual subcortical volume in each community and in the entire sample. In men, no case-control differences at the factor score level survived FDR correction, and only pallidum volume did for Community 1 (Table 3, Table S5). Importantly, the effect sizes of case-control differences within communities were larger than those of the whole subsample (Table 3, Table S4, and Table S5).

**Table 3:** Comparison of the mean of three factor scores between ADHD patients and controls in each community.

| Community | Basal ganglia |  |  |  | Limbic system |  |  |  | Thalamus |  |  |  |
| --- | --- | --- | --- | --- | --- | --- | --- | --- | --- | --- | --- | --- |
|  | Mean factor scores |  | P value | Cohen's d<br>(95% CIs) | Mean factor scores |  | P value | Cohen's d<br>(95% CIs) | Mean factor scores |  | P value | Cohen's d<br>(95% CIs) |
|  | Patients | Controls |  |  | Patients | Controls |  |  | Patients | Controls |  |  |
| **Boys** | -0.10<br>(0.90) | 0.13<br>(0.89) | **2.0e-04** | -0.26<br>(-0.38 - -0.13) | -0.01<br>(0.83) | 0.01<br>(0.84) | 0.74 | -0.02<br>(-0.15 - 0.10) | 0.02<br>(0.81) | -0.03<br>(0.82) | 0.40 | 0.07<br>(-0.06 - 0.19) |
| **1** | 0.44<br>(0.74) | 0.75<br>(0.71) | **3.0e-03** | -0.42<br>(-0.67 - -0.17) | 0.30<br>(0.72) | -0.18<br>(0.75) | **3.5e-06** | 0.65<br>(0.39 - 0.90) | -0.60<br>(0.67) | -0.51<br>(0.68) | 0.41 | -0.13<br>(-0.37 - 0.13) |
| **2** | -0.61<br>(0.64) | -0.75<br>(0.73) | 0.18 | 0.21<br>(-0.04 - 0.46) | -0.10<br>(0.64) | 0.14<br>(0.66) | **0.01** | -0.37<br>(-0.63 - -0.12) | 0.60<br>(0.65) | 0.37<br>(0.65) | **0.01** | 0.37<br>(0.11 - 0.62) |
| **3** | -0.68<br>(0.72) | -0.06<br>(0.66) | **2.6e-09** | -0.90<br>(-1.17 - -0.62) | 0.62<br>(0.77) | 0.70<br>(0.72) | 0.46 | -0.11<br>(-0.37 - 0.16) | -0.21<br>(0.63) | -0.40<br>(0.65) | 0.06 | 0.30<br>(0.04 - 0.57) |
| **4** | 0.49<br>(0.73) | 0.36<br>(0.70) | 0.22 | 0.18<br>(-0.07 - 0.43) | -0.65<br>(0.67) | -0.58<br>(0.70) | 0.46 | -0.10<br>(-0.35 - 0.15) | 0.06<br>(0.77) | 0.59<br>(0.71) | **5.0e-07** | -0.71<br>(-0.97 - -0.45) |
| Cohen's d effect sizes* |  |  |  |  |  |  |  |  |  |  |  |  |
| **Men<sup>a</sup>** | 0.02<br>(0.88) | -0.03<br>(0.93) | 0.56 | 0.06<br>(-0.10 - 0.22) | 0.02<br>(0.85) | -0.04<br>(0.86) | 0.45 | 0.08<br>(-0.08 - 0.24) | 0.03<br>(0.85) | -0.06<br>(0.88) | 0.40 | 0.10<br>(-0.06 - 0.26) |
| **1** | 0.74<br>(0.72) | 0.59<br>(0.77) | 0.32 | 0.20<br>(-0.06 - 0.46) | -0.15<br>(0.80) | 0.07<br>(0.83) | 0.28 | -0.27<br>(-0.53 - 0.00) | -0.48<br>(0.76) | -0.65<br>(0.77) | 0.28 | 0.22<br>(-0.04 - 0.49) |
| **2** | -0.70<br>(0.71) | -0.74<br>(0.77) | 0.72 | 0.06<br>(-0.26 - 0.37) | 0.26<br>(0.63) | 0.37<br>(0.74) | 0.45 | -0.16<br>(-0.48 - 0.15) | 0.27<br>(0.68) | 0.15<br>(0.59) | 0.40 | 0.19<br>(-0.13 - 0.51) |
| **3** | -0.20<br>(0.71) | -- | -- | -- | 0.69<br>(0.83) | -- | -- | -- | -0.34<br>(0.63) | -- | -- | -- |
| **4** | -0.07<br>(0.65) | -0.24<br>(0.66) | 0.28 | 0.26<br>(-0.04 - 0.56) | -0.42<br>(0.72) | -0.62<br>(0.71) | 0.28 | 0.27<br>(-0.02 - 0.58) | 0.66<br>(0.73) | 0.61<br>(0.70) | 0.71 | 0.07<br>(-0.23 - 0.37) |
| Cohen's d effect sizes* |  |  |  |  |  |  |  |  |  |  |  |  |

Note: FDR corrected p value. Significant difference in bold. 95% CIs: 95% Confidence intervals. \*Cohen's s effect sizes come from t-test that compared mean factor scores between ADHD patients and healthy controls in each subsample and community. <sup>a</sup> Community 3 is absent in men, because no healthy controls were presented.

### ADHD clinical profiles and comorbidities in communities

Among boys with ADHD, information on the severity of IA and HI symptoms was available for n=355 (63.0%) and n=358 (63.5%), respectively. This information was also available for 135 men with ADHD (32.8%). Neither total ADHD symptoms nor IA/HI symptom levels differed between communities in either boys or men (not shown).

For the analysis of comorbidities, we concentrated only on the presence or absence of common psychiatric comorbidities in ADHD, since the assessment of psychiatric comorbidities had been done using varied instruments across cohorts. Information was available for 311 (55.2%) boys with ADHD. Among them, 120 (38.6%) reported comorbid psychiatric disorders (Table S6). Anxiety and oppositional defiant disorder (ODD) were most frequently reported, occurring in 9.6% and 16.4%, respectively. There was neither a difference in the presence of (any) comorbidity between communities (*χ*^2^=0.98, *p*=0.81), nor were anxiety or ODD more frequently reported in one community compared to any other (anxiety: *χ*^2^=4.95, *p*=0.18; ODD: *χ*^2^=5.09, *p*=0.17). In men with ADHD, 205 (49.8%) had available information; among them, 113 (55.1%) reported comorbid psychiatric disorders (Table S7). Mood disorder and substance use disorder (SUD) were most frequently reported, occurring in 32.4% and 22.9% of men with ADHD, respectively. Presence of (any) comorbidity was more frequent in Community 1 and Community 4 than in the other two communities (*χ*^2^=15.63, *p*=0.001). Mood disorder and SUD were most frequent in Community 1 and Community 4 (mood disorder: χ2=9.35, *p*=0.02; SUD: χ2=23.08; *p*=2.0e-05).

## Discussion

In this study, we set out to investigate whether previously reported small effect sizes of case-control brain volume differences in ADHD might be explained by (structured) heterogeneity. Factor analysis of volumetric covariance indicated that the latent structure of subcortical volumes consists of basal ganglia, limbic system, and thalamus in male participants. Different latent factors seemed to underlie subcortical organization in females. Given sample sizes considerations, we concentrated all subsequent analyses on males. Among them, we discerned four distinct communities, one of which did not comprise any healthy adult males. In the subsample of boys, effect sizes of several case-control differences were larger within specific communities than in the total sample. The substructure of the brain volumes did not seem to have a behavioral correlate at the level of ADHD symptom severity, but men with ADHD in two communities more frequently reported the presence of comorbidities than those within the other two communities.

Similar factor structures of subcortical brain volumes existed in boys and men, regardless of ADHD status. The observed three-factor structure - basal ganglia, limbic system, and thalamus - is consistent with functional neuroanatomy and neurodevelopmental connections (22). Interestingly, factor structures differed between male and female participants, and also among females across the lifespan. Sex differences in subcortical brain volumes have consistently been reported in previous studies. Some studies reported larger volumes of amygdala, pallidum, and putamen in males (23, 24); Other studies observed larger hippocampus, caudate nucleus, and thalamus in females (25-27). However, this is the first paper to report on different correlations between subcortical structures in the two sexes. It is interesting to speculate, whether such differences in subcortical brain volume organization may be related to differences in ADHD presentation and comorbidity profiles between sexes.

Both boys and men could be separated into communities based on subcortical volume modularity. The community structure observed was similar in cases and controls, as has been observed also in cognitive investigations of ADHD (8, 13), providing further evidence that heterogeneity among individuals with ADHD is ‘nested’ in normal variation(13). In the present study, four communities were observed in boys with and without ADHD and in men with ADHD, while in healthy men, only three communities were present. It seemed like community structure in healthy men simplified from four to three communities, whilst patients retained a four-community distribution; this may be consistent with findings of delayed maturation in ADHD (11, 28, 29), but more research in longitudinal samples is clearly needed.

Effect sizes for case-control differences reported for subcortical volumes have always been small. The largest study of subcortical brain volumes in ADHD, performed by the ENIGMA-ADHD Working Group, reported effect sizes ranging from *d*=-0.19 to −0.10 across the lifespan, with largest effects in children (11). Case-control differences within each community showed that (a) not every community had significant differences for a specific volume, and (b) among those communities showing significant differences at the factor level, effect sizes ranged from *d*=-0.90 (95% CIs [−1.17, −0.62]) to 0.65 (95% CIs [0.39, 0.90]), which were considerably larger than the largest effect size observed in the full cohort, which was −0.26 (95% CIs [−0.38, −0.13]) (Table 3). Similar trends were also present for individual subcortical brain volumes (Tables S4 and S5). The current results highlight the neuroanatomical heterogeneity in the population and suggest that brain-based ADHD subtypes may exist.

As in the ENIGMA-ADHD and previous meta-analyses, case-control differences in the basal ganglia factor all pointed to smaller volumes in ADHD patients (11). More differentiated results were observed for the limbic system and thalamus. For the limbic system (and its components, amygdala and hippocampus), larger volumes were seen in boys with ADHD in Community 1, whereas the cases in Community 2 had smaller volumes. For the thalamus, we observed larger volumes in individuals with ADHD in Community 2 and Community 3, whereas those with ADHD in Community 4 had smaller volumes than healthy controls. Such findings may reconcile inconsistencies in the direction of effects reported for these structures in previous studies. Case-control differences were not significant in adult males. This result corroborates the earlier findings that developmental brain-structural differences observed with MRI in ADHD may normalize in adulthood (11, 28, 29).

To analyze the significance of the brain-structure-based communities for clinical presentation of ADHD, we explored potential differences between ADHD patients in the different communities. The communities did not appear to be associated with the severity of ADHD symptoms. This might have been a result of our limited sample size for these analyses, but several other neuropsychological studies had not found differences in ADHD symptom severity across subgroups either (13, 15). We did find some indication of clinical relevance of the communities when analyzing the presence of comorbidity: adult males with ADHD in Community 1 and Community 4 more frequently reported comorbidities than those in the other two communities, in particular mood disorder and SUD. Community 1 and Community 4 were characterized by relatively larger basal ganglia across the entire sample, which may be consistent with a previous study reporting increased basal ganglia volume in long-term substance abusers (30). The lack of significant associations with symptom severity and the limited findings for comorbidities may be due to insufficient power of the analyses in individual communities. Replication in independent samples with larger sample sizes is needed.

The strengths of the current study include the use of the large sample size of the ENIGMA-ADHD dataset to explore neuroanatomic subgroups, which provides us with the opportunity to better understand the small effect sizes of case-control differences in ADHD. A potential limitation is the arbitrariness of using the modularity algorithm; the application of different classification methodologies could result in different communities. However, in this study, we applied a widely-used technique and got a good approximation across subsamples. A second limitation was the heterogeneity of the ENIGMA-ADHD dataset, where several different diagnostic instruments had been used, and the fact that sample sizes dramatically shrank when we examined associations for single communities. Thirdly, we only focused on subcortical brain volumes in this study, as these have been most consistently associated with ADHD. However, differences between ADHD patients and controls are also observed in cortical measures, especially in surface area (28). Therefore, the clinical relevance of communities might be increased if taking into account cortical features. Lastly, since the factor structures differed between males and females, we only applied CD analyses in males, given sample size constraints in females. The neuroanatomic profiles of subcortical brain volumes in females and the heterogeneity among genders needs to be better understood.

To conclude, using subcortical MRI data from the ENIGMA-ADHD Working Group, we succeeded in stratifying our sample into neuroanatomically more homogeneous subgroups with preliminary links to the clinical presentation of ADHD. Our study may provide groundwork for future studies related to neuroanatomical heterogeneity in ADHD to increase our understanding of its neuro-pathophysiology.

## Acknowledgements

Ting Li is supported by China Scholarship Council (CSC) under the Grant CSC n° 201507720006. ENIGMA received funding from the National Institutes of Health (NIH) Consortium grant U54 EB020403, supported by a cross-NIH alliance that funds Big Data to Knowledge Centers of Excellence (BD2K). Support was also received from the European Community’s Horizon 2020 Programme (H2020/2014 – 2020) under grant agreements n° 667302 (CoCA) and n° 728018 (Eat2beNICE). Martine Hoogman and Barbara Franke were supported by personal grants from the Netherlands Organization for Scientific Research (NWO) Innovation Program (Veni grant 91619115 to MH; Vici grant 016-130-669 to BF). Lastly, we also gratefully acknowledge support from the European College for Neuropsychopharmacology (ECNP) for the ECNP Network ADHD across the Lifespan.

In addition to the people listed as co-authors, the following people are currently members of the ENIGMA-ADHD Working Group: Anatoly Anikin, Philip Asherson, Alexandr Baranov, Tiffany Chaim-Avanicini, Anders M. Dale, Alysa E. Doyle, Terry L. Jernigan, Sarah Hohmann, Dmitry Kapilushniy, Mitul Mehta, Leyla Namazova-Baranova, Stephanie E. Novotny, Eileen Oberwelland Weiss, Lena Schwarz, and Theo GM van Erp.

## Declaration of interests

Jan K Buitelaar has been in the past 3 years a consultant to / member of advisory board of / and/or speaker for Shire, Roche, Medice, and Servier. He is not an employee of any of these companies, and not a stock shareholder of any of these companies. He has no other financial or material support, including expert testimony, patents, royalties. In the past year, Dr. Faraone received income, potential income, travel expenses continuing education support and/or research support from Tris, Otsuka, Arbor, Ironshore, Shire, Akili Interactive Labs, Enzymotec, Sunovion, Supernus, and Genomind. With his institution, he has US patent US20130217707 A1 for the use of sodium-hydrogen exchange inhibitors in the treatment of ADHD. He also receives royalties from books published by Guilford Press: Straight Talk about Your Child’s Mental Health, Oxford University Press: Schizophrenia: The Facts, and Elsevier: ADHD: Non-Pharmacologic Interventions. He is Program Director of www.adhdinadults.com. Barbara Franke has received educational speaking fees from Medice. All other authors do not report possible conflicts of interests.

## Notes

http://enigma.ini.usc.edu/ongoing/enigma-adhd-working-group/

