## Supplementary for "Characterizing neuroanatomic heterogeneity in people with and without ADHD based on subcortical brain volumes"

**Table S1. Additional information of methods in each participating site.**

| **Sample** | **N** | **Age (SD)** | **N**  **missingness** | **Free-surfer version** | **Field strength** | **Classification for diagnosis** | **Instrument for symptom rating** | **Instrument for comorbidity assessment** | **IQ instrument** |
| --- | --- | --- | --- | --- | --- | --- | --- | --- | --- |
| **ADHD_Rubia** | 37 | 12.3 (1.1) | 5 | 5.3 | 3 Tesla | DSM-IV | SDQ for HI and Conners t-score for IA | Comorbid disorders were exclusion criteria | WASI |
| **ADHD_UKA** | 134 | 10.8 (1.3) | 31 | NA | NA | NA | German Parental and Teacher Report on ADHD | K-SADS-PL | NA/ CPM/WASI/WISC-IV full scale |
| **ADHD200_KKI** | 85 | 10.3 (1.4) | 9 | 5.3 | 1.5 Tesla | DSM-IV | Conners Parent Rating scale revised Long version (CPRSLV) | NA | WISC-IV |
| **ADHD200_NYU** | 182 | 10.5 (2.1) | 33 | 5.3 | 3 Tesla | DSM-IV | Conners Parent Rating scale revised Long version (CPRSLV) | NA | WASI |
| **ADHD200_OHSU** | 90 | 9.3 (1.3) | 23 | 5.3 | 3 Tesla | DSM-IV | Conners Parent Rating scale 3rd edition | NA | Block Design, Vocabulary and information subtests of WISC-IV |
| **ADHD200_Peking** | 213 | 11.4 (1.8) | 15 | 5.3 | 3 Tesla | DSM-IV | ADHD Rating Scale IV (ADHD-RS) | NA | WISCC-R |
| **Bergen_SVG** | 52 | 10.1 (1.3) | 2 | 5.3 | 3 Tesla | DSM-IV | K-SADS-PL | K-SADS-PL | WISC-IV |
| **CAPS_UZH** | 48 | 11.2 (1.6) | 6 | NA | NA | DSM-IV | Parent Conners | K-SADS-PL | NA |
| **DAT_London** | 19 | 13.1 (0.7) | 0 | 5.3 | 3 Tesla | DSM-IV | NA | NA | Vocabulary, Similarities, Picture Completion and Block Design, and information subtests of WISC-IV |
| **EPOD*** | 45/46 | 10.8 (0.9)  /28.1 (4.6) | 5/0 | 5.3 | 3 Tesla | DSM-IV | DBD-RS/ADHD-RS | DISC-IV/MINI-Plus | Vocabulary and Block Design of WISC-IIIR/DART |
| **NeuroImage_ADAM*** | 40/8 | 13.3 (0.9)  /23.1 (0.4) | 4/4 | 5.3 | 1.5 Tesla | DSM-IV | K-SADS-PL | K-SADS-PL | Vocabulary and Block Design, and information subtests of WISC-IV |
| **NeuroImage_NIJM*** | 40/6 | 12.6 (1.3)  /23.5 (0.5) | 6/0 | 5.3 | 1.5 Tesla | DSM-IV | K-SADS-PL | K-SADS-PL | Vocabulary and Block Design, and information subtests of WISC-IV |
| **NICAP** | 141 | 9.9 (0.5) | 28 | 5.3 |  |  | NA | DISC-IV | WASI vocabulary matrix reasoning |
| **NICHE** | 149 | 10.2 (1.7) | 2 | 5.1 | 1.5Tesla | DSM-IV | NA | DSM-IV | Vocabulary and Block Design, and information subtests of WISC-III |
| **UCHZ*** | 30/26 | 10.8 (1.6)  /35.0 (9.3) | 2/2 | 5.3 | 3 Tesla | NA | Parent Conners | KSADS-PL | HAWIK-IV |
| **ZI-CAPS** | 32 | 12.6 (1.3) | 1 | 5.3 | 3 Tesla | DSM-IV | Teachers Conners Kiddie SADS | ODD and CD with structured clinical interview | Subscales of HAWIK-IV |
| **UAB-ADHD*** | 57/115 | 10.2 (2.5)  /35.2 (8.1) | 0/0 | 5.3 | 3 Tesla | DSM-IV | NA | NA | Subscales of HAWIK-IV/ WISC |
| **ADHD WUE** | 104 | 40.8 (11.0) | 5 | 5.3 | 1.5Tesla | DSM-IV | DSM-IV interview | SCID1 | MWT-B |
| **ADHD-DUB1** | 21 | 27.1 (6.7) | 5 | 5.3 | 3Tesla | DSM-IV | Conners Adults ADHD rating scale observer | SCID1 | Verbal comprehension, perceptual, reasoning, working memory and processing speed subtests of WAIS-IV |
| **ADHD-DUB2** | 17 | 35.3 (10.1) | 0 | 5.3 | 1.5Tesla | DSM-IV | Conners Adults ADHD rating scale observer | SCID1 | NA |
| **ADHD-Mattos** | 21 | 25.3 (2.7) | 0 | 5.1 | 3Tesla | DSM-IV | K-SADS adapted for adults | MINI | WASI |
| **Bergen_adultADHD** | 89 | 32.1 (6.4) | 0 | 5.3 | 3Tesla | DSM-IV/ICD-10 | NA | NA | WASI |
| **IMpACT_NL** | 212 | 37.5 (10.6) | 4 | 5.3 | 1.5Tesla | DSM-IV | DSM-IV interview | SCID1&2 | Vocabulary and block design subtests of WASI |
| **MGH_ADHD** | 122 | 38.9 (10.7) | 2 | 5.1 | 1.5Tesla | DSM-IV | DSM-IV interview | SCID1 | Vocabulary and block design subtests of WASI |
| **MTA** | 121 | 24.8 (1.2) | 0 | 5.3 | 3Tesla | DSM-IV | NA | NA | WISC-III |
| **NYU-ADHD** | 61 | 35.0 (8.3) | 0 | 5.3 | 3Tesla | DSM-IV | NA | SCID1 | WASI |
| **SAOPAULO** | 115 | 29.1 (4.9) | 0 | 5.3 | NA | NA | NA | SCID1 | WASI |
| **Tuebingen** | 16 | 30.9 (6.1) | 0 | NA | NA | NA | NA | NA | NA |

*Note: * Cohorts included both children and adult participants.*

**Table S2. The distribution of boys (patients/controls) in each community**

| **Sample** | **Patients-Community (%)** | | | |  | **Controls-Community (%)** | | | |
| --- | --- | --- | --- | --- | --- | --- | --- | --- | --- |
|  | **1** | **2** | **3** | **4** |  | **1** | **2** | **3** | **4** |
| **ADHD_Rubia*** | 6 (27.2%) | 0 | 16 (72.7%) | 0 |  | 5 (33.3%) | 0 | 10 (66.7%) | 0 |
| **ADHD_UKA*** | 3 (3.6%) | 54 (64.3%) | 23 (27.4%) | 4 (4.8%) |  | 1 (4.8%) | 6 (28.6%) | 13 (61.9%) | 1 (4.8%) |
| **ADHD200_KKI*** | 3 (21.4%) | 4 (28.6%) | 0 | 7 (50.0%) |  | 9 (23.1%) | 2 (5.1%) | 1 (2.6%) | 27 (69.2%) |
| **ADHD200_NYU*** | 19 (24.7%) | 5 (6.5%) | 7 (9.1%) | 46 (59.7%) |  | 9 (25.7%) | 0 | 1 (2.7%) | 25 (71.4%) |
| **ADHD200_OHSU** | 3 (15.8%) | 8 (42.1%) | 4 (21.1%) | 4 (21.1%) |  | 8 (28.6%) | 15 (53.6%) | 4 (14.3%) | 1 (3.6%) |
| **ADHD200_Peking** | 18 (25.4%) | 21 (29.6%) | 14 (19.7%) | 18 (25.4%) |  | 25 (33.3%) | 20 (26.7%) | 16 (21.3%) | 14 (18.7%) |
| **Bergen_SVG** | 9 (47.4%) | 0 (0%) | 9 (47.4%) | 1 (5.3%) |  | 6 (30.0%) | 2 (10.0%) | 12 (60.0%) | 0 |
| **CAPS_UZH** | 4 (21.1%) | 3 (15.8%) | 12 (63.2%) | 0 |  | 3 (21.4%) | 2 (14.3%) | 9 (64.3%) | 0 |
| **DAT_London** | 0 | 5 (45.5%) | 6 (54.5%) | 0 |  | 0 | 8 (100.0%) | 0 | 0 |
| **EPOD** | 1 (2.2%) | 25 (55.6%) | 18 (40.0%) | 1 (2.2%) |  | - | - | - | - |
| **NeuroImage_ADAM** | 0 | 1 (12.5%) | 0 | 7 (87.5%) |  | 9 (39.1%) | 2 (8.7%) | 1 (4.3%) | 11 (47.8%) |
| **NeuroImage_NIJM** | 6 (40.0%) | 0 | 0 | 9 (60.0%) |  | 0 | 0 | 0 | 6 (100.0%) |
| **NICAP** | 8 (16%) | 17 (34.0%) | 10 (20.0%) | 15 (30.0%) |  | 8 (17.8%) | 14 (31.1%) | 17 (37.8%) | 6 (13.3%) |
| **NICHE** | 6 (9.5%) | 25 (39.7%) | 8 (12.7%) | 24 (38.1%) |  | 5 (7.8%) | 25 (39.1%) | 9 (14.1%) | 25 (39.1%) |
| **UAB-ADHD** | 10 (43.5%) | 2 (8.7%) | 2 (8.7%) | 9 (39.1%) |  | 12 (60.0%) | 0 | 5 (25.0%) | 3 (15.0%) |
| **UCHZ** | 6 (85.7%) | 0 | 0 | 1 (14.3%) |  | 6 (60.0%) | 0 | 1 (10.0%) | 3 (30%) |
| **ZI-CAPS** | 8 (50.0%) | 4 (25.0%) | 4 (25.5%) | 0 |  | 4 (66.7%) | 0 | 2 (33.3%) | 0 |
| **Total** | **110 (19.5%)** | **174 (30.9%)** | **133 (23.6%)** | **146 (25.9%)** |  | **110 (25.6%)** | **96 (22.4%)** | **101 (23.5%)** | **122 (28.4%)** |

*Note: The distribution of participants based on factor scores derived from EFA on boys (with and without ADHD). * Cohorts involved in leave-one-out analyses.*

**Table S3. The distribution of adult men (patients/controls) in each community**

| **Sample** | **Patients-Community (%)** | | | |  | **Controls-Community (%)** | | | |
| --- | --- | --- | --- | --- | --- | --- | --- | --- | --- |
|  | **1** | **2** | **3** | **4** |  | **1** | **2** | **3** | **4** |
| **ADHD-DUB1** | 4 (57.1%) | 0 | 1 (14.3%) | 2 (28.6%) |  | 4 (66.7%) | 0 | 2 (33.3%) | 0 |
| **ADHD-DUB2** | 0 | 6 (42.9%) | 3 (21.4%) | 5 (35.7%) |  | - | - | - | - |
| **ADHD-Mattos** | 8 (57.1%) | 2 (14.3%) | 1 (7.1%) | 3 (21.4%) |  | - | - | - | - |
| **ADHD WUE** | 1 (3.7%) | 15 (55.6%) | 4 (14.8%) | 7 (25.9%) |  | 0 | 15 (62.5%) | 5 (20.8%) | 4 (16.7%) |
| **Bergen_adultADHD** | 14 (58.3%) | 1 (4.2%) | 9 (37.5%) | 0 |  | 9 (47.4%) | 1 (5.3%) | 9 (47.4%) | 0 |
| **EPOD** | 9 (19.6%) | 21 (45.7%) | 12 (26.1%) | 4 (8.7%) |  | - | - | - | - |
| **IMpACT_NL** | 2 (4.5%) | 7 (15.9%) | 0 | 35 (79.5%) |  | 4 (9.5%) | 5 (11.9%) | 0 | 33 (78.6%) |
| **MGH_ADHD** | 34 (97.1%) | 0 | 1 (2.9%) | 0 |  | 26 (100%) | 0 | 0 | 0 |
| **MTA** | 21 (30%) | 8 (11.4%) | 8 (11.4%) | 33 (47.1%) |  | 8 (28.6%) | 6 (21.4%) | 2 (7.1%) | 12 (42.9%) |
| **NeuroImage_ADAM** | 4 (80%) | 0 | 0 | 1 (20%) |  | - | - | - | - |
| **NeuroImage_NIJM** | 2 (50%) | 1 (25%) | 0 | 1 (25%) |  | - | - | - | - |
| **NYU-ADHD** | 10 (62.5%) | 0 | 6 (37.5%) | 0 |  | 14 (77.8%) | 0 | 4 (22.2%) | 0 |
| **SAOPAULO** | 5 (10.9%) | 14 (30.4%) | 6 (13.0%) | 21 (45.7%) |  | 5 (14.7%) | 13 (38.2%) | 3 (8.8%) | 13 (38.3%) |
| **Tuebingen** | 3 (23.1%) | 7 (53.8%) | 3 (23.1%) | 0 |  | - | - | - | - |
| **UAB-ADHD** | 8 (20%) | 20 (50.0%) | 7 (17.5%) | 5 (12.5%) |  | 4 (10.8%) | 21 (56.8%) | 9 (24.3%) | 3 (8.1%) |
| **UCHZ** | 2 (28.6%) | 1 (14.3%) | 1 (14.3%) | 3 (42.9%) |  | 0 | 2 (28.6%) | 1 (14.3%) | 4 (57.1%) |
| **Total** | **127 (30.8%)** | **103 (25.0%)** | **62 (15.0%)** | **120 (29.1%)** |  | **74 (30.7%)** | **63 (26.1%)** | **35 (14.5%)** | **69 (28.6%)** |

*Note: The distribution of participants based on factor scores derived from derived from EFA on men (with and without ADHD).*

**Table S4. Comparison of subcortical brain volumes of each community in subsample of boys**

|  | **Total sample** | | | **Community 1** | | | **Community 2** | | |
| --- | --- | --- | --- | --- | --- | --- | --- | --- | --- |
|  | **Mean volume**  **Patients/Controls** | ***P* value** | **Cohen’s d**  **(95% CIs)** | **Mean volume**  **Patients/Controls** | ***P* value** | **Cohen’s d**  **(95% CIs)** | **Mean volume**  **Patients/Controls** | ***P* value** | **Cohen’s d**  **(95% CIs)** |
| **Accumbens** | 683.5 (112.5)/  702.6 (118.2) | **0.03** | -0.16  (-0.29 - -0.03) | 750.7 (112.7)/  760.2 (111.4) | 0.70 | 0.05  (-0.20 - 0.30) | 616.6 (90.2)/  625.0 (102.5) | 0.39 | -0.12  (-0.38 - 0.13) |
| **Caudate** | 4117.4 (561.2)/  4217.7 (550.7) | **0.04** | -0.15  (-0.28 - -0.03) | 4088.1 (488.0)/  4351.3 (538.6) | 0.07 | -0.26  (-0.51 - -0.01) | 4142.0 (582.1)/  3983.3 (572.7) | 0.16 | 0.20  (-0.05 - 0.46) |
| **Putamen** | 6274.0 (714.0)/  6448.5 (735.3) | **8.8e-04** | -0.24  (-0.36 - -0.11) | 6645.4 (593.9)/  6993.3 (615.3) | **0.01** | -0.36  (-0.61 - -0.11) | 5897.4 (575.4)/  5787.7 (603.9) | 0.48 | 0.10  (-0.15 - 0.35) |
| **Pallidum** | 1840.5 (247.8)/  1880.8 (259.7) | **0.05** | -0.14  (-0.27 - -0.01) | 1805.5 (201.1)/  1971.7 (206.5) | **1.2e-05** | -0.63  (-0.89 - -0.38) | 1843.1 (218.8)/  1721.4 (250.2) | **5.0e-04** | 0.51  (0.25 - 0.76) |
| **Amygdala** | 1633.5 (202.4)/  1668.1 (216.5) | **0.03** | -0.15  (-0.28 - -0.03) | 1687.2 (179.7)/  1665.2 (209.2) | **0.002** | 0.45  (0.20 - 0.70) | 1613.9 (188.3)/  1638.4 (184.9) | **0.01** | -0.38  (-0.64 - -0.13) |
| **Hippocampus** | 4284.3 (476.6)/  4249.8 (447.8) | **0.05** | 0.14  (0.02 - 0.27) | 4214.9 (434.1)/  4131.8 (402.5) | **1.0e-04** | 0.57  (0.31 - 0.82) | 4427.3 (437.1)/  4367.3 (422.7) | 0.84 | -0.03  (-0.28 - 0.23) |
| **Thalamus** | 7867.0 (854.7)/  7871.6 (733.5) | 0.39 | 0.06  (-0.06 - 0.19) | 7327.8 (709.6)/  7608.0 (707.7) | 0.68 | -0.05  (-0.30 - 0.20) | 8372.5 (782.1)/  8116.7 (661.1) | 0.15 | 0.21  (-0.04 - 0.46) |

Continued

|  | **Community 3** | | | **Community 4** | | |
| --- | --- | --- | --- | --- | --- | --- |
|  | **Mean volume**  **Patients/Controls** | ***P* value** | **Cohen’s d**  **(95% CIs)** | **Mean volume**  **Patients/Controls** | ***P* value** | **Cohen’s d**  **(95% CIs)** |
| **Accumbens** | 673.3 (95.2)/  731.3 (111.4) | **1.8e-05** | -0.65  (-0.93 - -0.38) | 712.2 (106.3)/  672.4 (97.4) | **0.01** | 0.38  (0.23 - 0.63) |
| **Caudate** | 3870.7 (530.7)/  4149.2 (487.7) | **0.001** | -0.49  (-0.76 - -0.22) | 4310.5 (541.5)/  4335.8 (527.5) | 0.43 | -0.11  (-0.36 - 0.14) |
| **Putamen** | 5941.7 (671.8)/  6339.7 (609.0) | **1.0e-05** | -0.69  (-0.96 - -0.42) | 6661.0 (616.1)/  6479.7 (539.1) | **0.04** | 0.30  (0.04 - 0.55) |
| **Pallidum** | 1640.3 (199.7)/  1756.1 (212.7) | **3.3e-04** | -0.55  (-0.82 - -0.28) | 2024.4 (213.5)/  2049.6 (221.6) | 0.20 | -0.19  (-0.44 - 0.07) |
| **Amygdala** | 1727.7 (210.9)/  1809.5 (208.7) | **0.03** | -0.35  (-0.62 - -0.09) | 1537.9 (180.8)/  1555.6 (182.7) | 0.24 | -0.17  (-0.42 - 0.08) |
| **Hippocampus** | 4504.2 (465.9)/  4494.1 (410.6) | 0.10 | 0.25  (-0.02 - 0.51) | 4007.2 (408.1)/  4042.0 (415.9) | 0.27 | -0.16  (-0.41 - 0.09) |
| **Thalamus** | 7694.1 (809.3)/  7683.2 (701.6) | **0.03** | 0.34  (0.08 - 0.61) | 7873.7 (761.4)/  8172.7 (684.4) | **1.0e-05** | -0.68  (-0.94 - -0.42) |

*Note: FDR corrected p value. Significant difference in bold.95% CIs: 95% Confidence intervals.* *Adjusted mean volume estimate and SD of the mean corrected for age, age^2, ICV and sample sites.*

**Table S5. Comparison of subcortical brain volumes of each community in subsample of adult men**

|  | **Total sample** | | | **Community 1** | | |
| --- | --- | --- | --- | --- | --- | --- |
|  | **Mean volume**  **Patients/Controls** | ***P* value** | **Cohen’s d**  **(95% CIs)** | **Mean volume**  **Patients/Controls** | ***P* value** | **Cohen’s d**  **(95% CIs)** |
| **Accumbens** | 636.9 (127.8)/  647.7 (144.7) | 0.74 | -0.04  (-0.20 - 0.12) | 736.2 (124.6)/  757.5 (120.2) | 0.62 | -0.16  (-0.42 - 0.10) |
| **Caudate** | 3972.9 (528.9)/  3922.0 (520.7) | 0.55 | 0.11  (-0.04 - 0.27) | 3999.0 (602.2)/  3962.1 (525.7) | 0.74 | 0.06  (-0.21 - 0.32) |
| **Putamen** | 5910.3 (733.7)/  5890.6 (732.9) | 0.74 | 0.07  (-0.09 - 0.23) | 6434.4 (655.3)/  6291.6 (681.2) | 0.4 | 0.25  (-0.01 - 0.52) |
| **Pallidum** | 1760.8 (262.5)/  1738.1 (233.8) | 0.55 | 0.11  (-0.05 - 0.27) | 1831.5 (253.7)/  1735.8 (200.8) | **0.03** | 0.43  (0.16 - 0.69) |
| **Amygdala** | 1631.9 (200.3)/  1629.0 (210.9) | 0.74 | 0.04  (-0.12 - 0.20) | 1643.4 (190.2)/  1703.7 (204.0) | **0.17** | -0.34  (-0.60 - -0.07) |
| **Hippocampus** | 4383.0 (464.6)/  4226.8 (501.0) | 0.55 | 0.12  (-0.04 - 0.28) | 4192.7 (463.0)/  4204.3 (522.7) | 0.79 | -0.04  (-0.30 - 0.22) |
| **Thalamus** | 8121.6 (870.3)/  8059.9 (919.3) | 0.74 | 0.04  (-0.12 - 0.20) | 7617.4 (790.3)/  7560.4 (823.2) | 0.84 | 0.03  (-0.24 - 0.29) |

Continued

|  | **Community 2** | | | **Community 4** | | |
| --- | --- | --- | --- | --- | --- | --- |
|  | **Mean volume**  **Patients/Controls** | ***P* value** | **Cohen’s d**  **(95% CIs)** | **Mean volume**  **Patients/Controls** | ***P* value** | **Cohen’s d**  **(95% CIs)** |
| **Accumbens** | 553.9 (92.8)/  553.7 (104.9) | 0.74 | 0.11  (-0.21 - 0.42) | 583.5 (86.9)/  580.6 (97.5) | 0.62 | 0.16  (-0.13 - 0.47) |
| **Caudate** | 3845.2 (494.1)/  3796.9 (541.7) | 0.74 | 0.13  (-0.19 - 0.44) | 4021.7 (476.0)/  3989.6 (474.8) | 0.74 | 0.10  (-0.20 - 0.41) |
| **Putamen** | 5406.1 (563.0)/  5507.8 (646.0) | 0.74 | -0.07  (-0.39 - 0.24) | 5790.8 (596.4)/  5686.2 (590.3) | 0.48 | 0.26  (-0.04 - 0.56) |
| **Pallidum** | 1654.9 (244.0)/  1610.1 (221.9) | 0.52 | 0.26  (-0.06 - 0.57) | 1879.1 (240.6)/  1871.2 (219.5) | 0.74 | 0.07  (-0.23 - 0.37) |
| **Amygdala** | 1643.5 (159.4)/  1676.5 (163.5) | 0.62 | -0.18  (-0.50 - 0.13) | 1508.7 (161.2)/  1470.5 (178.1) | 0.41 | 0.30  (0.00 - 0.61) |
| **Hippocampus** | 4603.9 (339.0)/  4621.5 (444.4) | 0.74 | -0.06  (-0.38 - 0.25) | 4246.5 (427.4)/  4208.9 (392.1) | 0.74 | 0.12  (-0.18 - 0.42) |
| **Thalamus** | 8399.8 (765.2)/  8286.6 (698.3) | 0.74 | 0.11  (-0.20 - 0.43) | 8637.2 (749.6)/  7568.1 (890.0) | 0.74 | 0.09  (-0.21 - 0.39) |

*Note: FDR corrected p value. 95% CIs: 95% Confidence intervals.* *Adjusted mean volume estimate and SD corrected for age, age^2, ICV and sample sites.*

**Table S6. ADHD comorbidities in each cohort in boys with ADHD**

| **Sample*** | **Available/missing** | **Comorbidities**  **(yes/no)** | **Anxiety**  **(yes/no)** | **ODD**  **(yes/no)** |
| --- | --- | --- | --- | --- |
| **ADHD_Rubia** | 22/0 | 0/22 | 0/22 | 0/22 |
| **ADHD_UKA** | 83/1 | 29/54 | 6/77 | 10/73 |
| **Bergen_SVG** | 19/0 | 15/4 | 6/13 | 10/9 |
| **CAPS_UZH** | 17/2 | 2/15 | 0/17 | 0/17 |
| **DAT_London** | 11/0 | 0/11 | - | - |
| **NeuroImage_ADAM** | 8/0 | 3/5 | - | - |
| **NeuroImage_NIJM** | 15/0 | 9/6 | - | - |
| **NICAP** | 50/0 | 21/29 | 13/37 | 7/43 |
| **NICHE** | 63/0 | 32/31 | 0/63 | 22/41 |
| **UCHZ** | 7/0 | 0/7 | 0/7 | 0/7 |
| **ZI-CAPS** | 16/0 | 1/15 | - | 1/15 |
| **Total** | 311/3 | 120/191 | 25/236 | 50/254 |

**No available data in ADHD200_KKI, ADHD200_NYU, ADHD200_OHSU, ADHD200_Peking, EPOD, and NICHE*

**Table S7. ADHD comorbidities in each cohort in adult men with ADHD**

| **Sample*** | **Available/missing** | **Comorbidities**  **(yes/no)** | **Mood disorder**  **(yes/no)** | **SUD**  **(yes/no)** |
| --- | --- | --- | --- | --- |
| **ADHD_DUB1** | 7/0 | 5/2 | 1/6 | 4/2 |
| **ADHD_DUB2** | 14/0 | 4/10 | 4/10 | 0/14 |
| **ADHD_Mattos** | 14/0 | 6/8 | 1/13 | 2/12 |
| **ADHD_WUE** | 22/5 | 17/5 | 8/13 | 4/17 |
| **IMpACT_NL** | 44/0 | 33/11 | 15/21 | 6/23 |
| **MGH_ADHD** | 35/0 | 32/3 | 20/15 | 22/13 |
| **NeuroImage_ADAM** | 3/2 | 0/3 | - | - |
| **NeuroImage_NIJM** | 4/0 | 1/3 | 1/0 | - |
| **NYU ADHD** | 14/2 | 7/7 | 3/9 | 2/9 |
| **SAOPAULO** | 46/0 | 8/38 | 8/38 | 0/46 |
| **UCHZ** | 2/5 | 0/2 | 0/2 | 0/2 |
| **Total** | 205/14 | 113/92 | 61/127 | 40/138 |

**No available data in ADHD_DUB1, ADHD_DUB2, ADHD_Mattos, IMpACT_NL, MGH_ADHD, Tuebingen, UAB-ADHD, and UCHZ. SUD: substance use disorder*

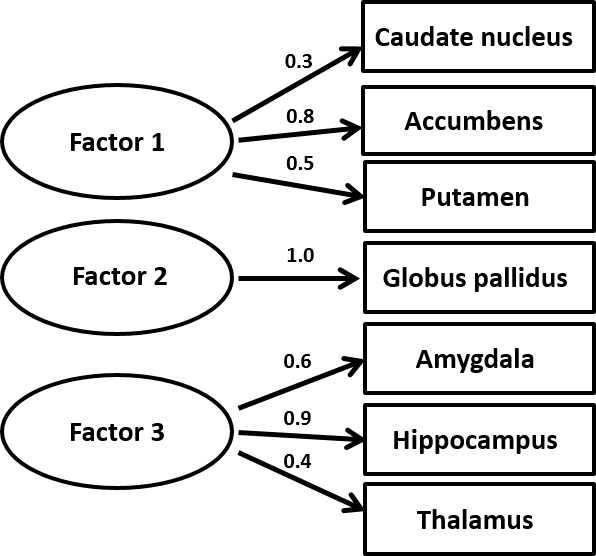

**Figure S1.** The three-factor model that was generated by EFA in girls with estimated factor loadings of the latent factors.

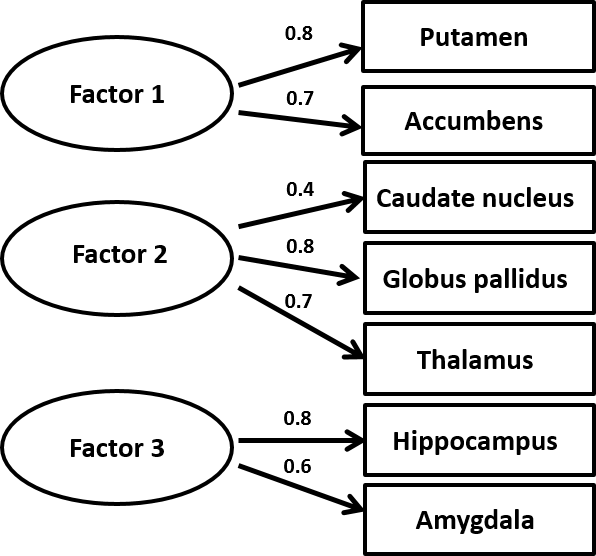

**Figure S2.** The three-factor model that was generated by EFA in adult women with estimated factor loadings of the latent factors.

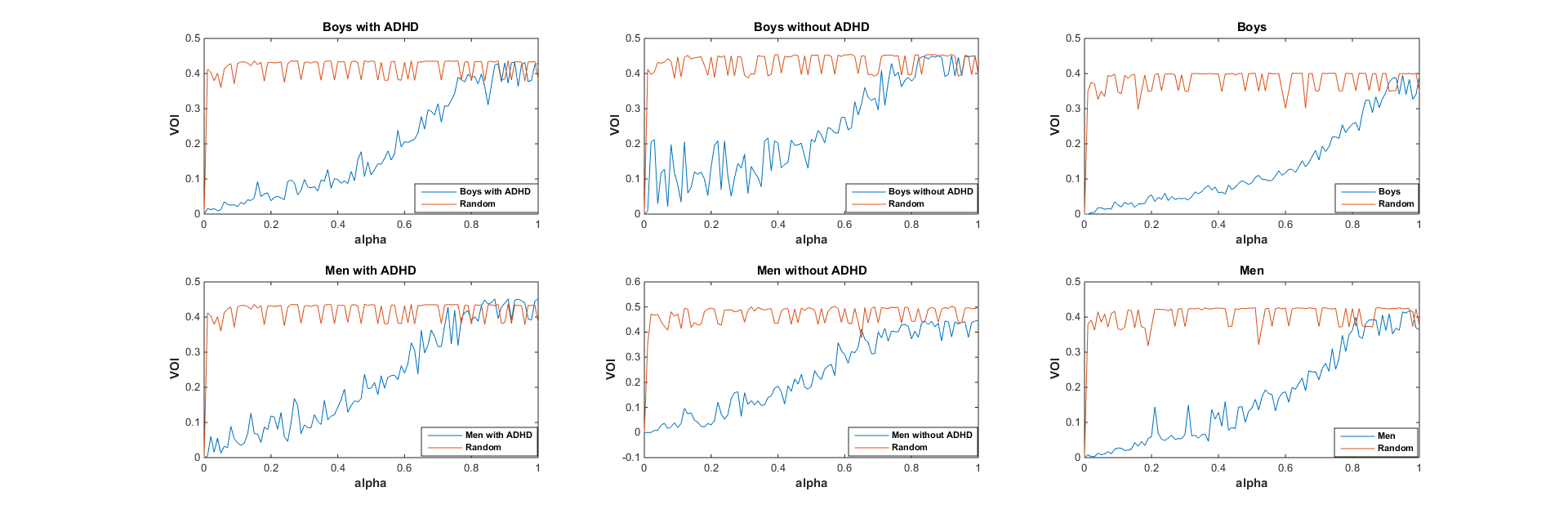

**Figure S3:** VOI in each subsample.
